# Neural Correlates of Value-Driven Spatial Attention

**DOI:** 10.1101/2021.06.28.450234

**Authors:** Ming-Ray Liao, Andy J. Kim, Brian A. Anderson

**Affiliations:** Texas A&M University, Department of Psychological and Brain Sciences, 4235 TAMU, College Station, TX 77843-4235

**Keywords:** spatial attention, reward, learning, eye movements

## Abstract

Reward learning has been shown to habitually guide spatial attention to regions of a scene. However, the neural mechanisms that support this bias in spatial orienting are unknown. In the present study, participants learned to orient to a particular quadrant of a scene (high-value quadrant) to maximize monetary gains. This learning was scene-specific, with the high-value quadrant varying across different scenes. During a subsequent test phase, participants were faster at identifying a target if it appeared in the high-value quadrant (valid), and initial saccades were more likely to be made to the high-value quadrant. fMRI analyses during the test phase revealed learning-dependent priority signals in the bilateral caudate tail and superior colliculus, frontal eye field, substantia nigra, and insula, paralleling findings concerning feature-based value-driven attention. In addition, ventral regions typically associated with scene selective and spatial information processing, including the hippocampus, parahippocampal gyrus, and temporo-occipital cortex, were also implicated. Taken together, our findings offer new insights into the neural architecture subserving value-driven attention, both extending our understanding of nodes in the attention network previously implicated in feature-based value-driven attention and identifying a ventral network of brain regions implicated in reward’s influence on scene-dependent spatial orienting.

## INTRODUCTION

Selectively representing relevant and important information is necessary for us to effectively navigate dense visual environments. Our brains must accordingly facilitate this process, and much evidence suggests that our perceptual experience is the product of a competitive process in which attention biases perception in favor of selected stimuli (Desimone & Duncan, 1995). Sources of attentional bias include top-down factors such as task goals (Wolfe et al., 1989; Folk et al. 1992), bottom-up factors such as physical salience (Theeuwes, 1992), and selection history from associative reward learning (Anderson et al., 2011; Anderson, 2016; Hickey et al., 2010), statistical regularities (Britton & Anderson, 2020; Wang & Theeuwes, 2018a, 2018b, 2018c), and aversive conditioning (Anderson & Britton, 2019; Nissens et al., 2017; Schmidt et al., 2015).

The role of associative reward learning in guiding feature-based attention is well established (see Anderson, 2016, for a review). Numerous studies have demonstrated that previously rewarding stimuli continue to bias attention into extinction (Anderson & Yantis, 2013; Liao & Anderson, 2020a; Milner et al., 2020), even when the reward-associated feature is nonsalient, no longer task-relevant, and no longer predictive of reward. The features commonly employed include but are not limited to colors (Anderson et al., 2011; Le Pelley et al., 2015), shapes (Della Libera & Chelazzi, 2009; Della Libera et al., 2011), orientations (Laurent et al., 2015), and object categories (Donohue et al., 2016; Hickey et al., 2015). The neural correlates of visual cortex value-driven attention for features are also well established and include the ventral visual cortex, frontal eye field, and caudate tail (Anderson et al., 2014; Anderson et al., 2016; Barbaro et al., 2017; Donohue et al., 2016; Hickey & Peelen, 2015, 2017; Kim & Anderson 2020a, 2020b; Kim & Hikosaka, 2013; Yamamoto et al., 2013; see Anderson, 2019, for a review). The early visual cortex (Itthipuripat et al., 2019; MacLean & Giesbrecht, 2015; Serences, 2008; Serences & Saproo, 2010) and insula (Wang et al., 2015) have also been implicated.

More recently, some studies have demonstrated that reward learning can also guide spatial attention (Anderson & Kim, 2018a, 2018b; Chelazzi et al., 2014; see also Liao & Anderson, 2020b). In Anderson and Kim (2018a), participants learned to associate a region in space within distinctive object-rich scenes with reward. After this learning, they performed a visual search task superimposed on the scenes. It was found that target identification was facilitated when the target appeared in the previously reward-associated region (valid trial). Eye movements were likewise biased towards the previously reward-associated region (Anderson & Kim, 2018a, 2018b). The neural mechanisms that support such an influence of reward on spatial attention remain to be investigated.

Evidence from both behavior (e.g., Anderson & Kim, 2019a, 2019b; Anderson & Yantis, 2012; Kim & Anderson, 2019a) and neuroimaging (e.g., Anderson et al., 2014, 2016; Anderson, 2017, 2019; Kim & Anderson, 2020a, 2020b) in the case of value-driven feature-based attention, and behavior in the case of value-driven spatial attention (Anderson & Kim, 2018a, 2018b; Liao & Anderson 2020b), suggest a common influence on the oculomotor system. This opens up the possibility of a common network of brain regions subserving both modes of value-modulated orienting. This includes the caudate tail, which is causally linked to eye movements (Yamamoto et al., 2012), along with the superior colliculus (to which the caudate tail projects via the substantia nigra pars reticulata; Yamamoto et al., 2012) and frontal eye field (to which the superior colliculus projects via the mediodorsal thalamus; Sommer & Wurtz, 2004).

A unique element of value-driven spatial attention is the reliance on object-rich scenes that can provide contextual information about where to guide attention (Brockmole & Henderson, 2006a, 2006b). With the inclusion of complex objects and spatial layout that collectively serve as a cue for a high-value region, we expect parts of the medial temporal lobe like the hippocampus and parahippocampal gyrus―not previously implicated in value-driven feature-based attention (see Anderson, 2019)―to play an important role in signaling scene-specific spatial biases. The caudate tail receives input via the ventral visual stream and in particular the visual cortico-striatal loop (Anderson, 2019; Seger, 2013); the ventral visual cortex is robustly activated by complex scenes in a manner modulated by reward (Barbaro et al., 2017; Hickey & Peelen, 2015, 2017), and the caudate tail runs adjacent to the hippocampus and surrounding parahippocampal gyrus, which play a well-defined role in spatial memory (Epstein & Kanwisher, 1998; Maguire et al., 1996; O’keefe & Nadel, 1978). Value-driven attention for low-level features are sensitive to specific scene contexts (Anderson, 2015; see also Gregoire et al., in press) which, along with the caudate tail’s proximity to the medial temporal lobe (Seger, 2013) and its connections with the superior colliculus (Yamamoto et al., 2012), raise the possibility that this network of brain regions is collectively involved in representing value-driven spatial attention.

Using human functional magnetic resonance imaging (fMRI), we employed a whole-brain approach to investigate the representation of task-irrelevant, value-driven spatial biases using the paradigm established by Anderson and Kim (2018a). Given the aforementioned considerations, we hypothesized that valid trials, in which the target of visual search requires an eye movement to the region of a scene previously associated with high value, would be associated with more robust activation (biased competition) in oculomotor regions of the brain previously implicated in value-driven feature-based attention (caudate tail, superior colliculus, frontal eye field) in addition to the hippocampus and parahippocampal gyrus given the reliance on scene context.

## METHODS

### Participants

Forty-seven participants (18-35 years of age, M = 22.83 years, SD = 4.55; 24 females, 23 males) were recruited from the Texas A&M Community. The demographic information for one participant was lost due to experimenter error. Participants were compensated with money earned in the experimental task. All reported normal or corrected-to-normal visual acuity and normal color vision. All procedures were approved by the Texas A&M University Institutional Review Board and conformed with the principles outlined in the Declaration of Helsinki. All participants provided written informed consent. Of the 47 recruited participants, 12 did not meet the required task performance to continue in the scanner (failed to robustly learn the pairings between locations in scenes and reward or could not perform that test phase task with sufficient accuracy), and one withdrew partway through the scan. The final sample consisted of 34 participants who completed the entire experiment, for which 33 of their demographic data is available (M = 22.33 years, SD = 4.36; 15 females, 18 males). The obtained sample size provided power (1-β) > 0.9 to replicate an effect of reward learning on eye movements in the test phase Anderson and Kim (2018a, 2018b) (computed using G*Power 3.1), and was similar to (and in most cases exceeded) the sample sizes used in prior studies of the neural correlates of value-driven attention (Anderson et al., 2014; Anderson, 2017; Itthipuripat et al., 2019; Barbaro et al., 2017; Hickey & Peelen, 2015; Kim & Anderson, 2019b, 2019c, 2020a, 2020b).

### Apparatus

In-lab tasks were completed on a Dell OptiPlex equipped with Matlab software and Psychophysics Toolbox extensions (Brainard, 1997). Stimuli were presented on a Dell P2717H Monitor. Participants viewed the monitor from a distance of approximately 70cm in a dimly lit room. Manual responses were entered using a standard keyboard. Eye-tracking was conducted using the EyeLink 1000 Plus system while head position was maintained using a manufacturer-provided chin rest (SR Research Ltd). Stimulus presentation during the fMRI portion was controlled by an Invivo SensaVue display system. The eye-to-screen distance was approximately 125cm. Responses were entered using Cedrus Lumina two-button response pads. An EyeLink 1000 Plus system was again used to track eye position.

### Training phase

Each trial began with a fixation cross (1.1º visual angle) that remained at the center of the screen until eye position had been registered within 1.8º of the fixation cross for a continuous period of 500ms (Figure 1). After which, a scene image was displayed that filled the entire computer screen. Four grey rectangular outlines (9.1º x 6.9º) were also displayed at the center of each quadrant, the center of which were 11.4º away from the center of the screen. The scene and rectangles remained on the screen until eye position had been registered within the boundary of one of the rectangles for a continuous period of 1000ms. After a 500ms blank screen, the reward feedback display was presented for 1500ms and consisted of the money earned on the current trial along with the updated total earnings.

**Figure 1.**
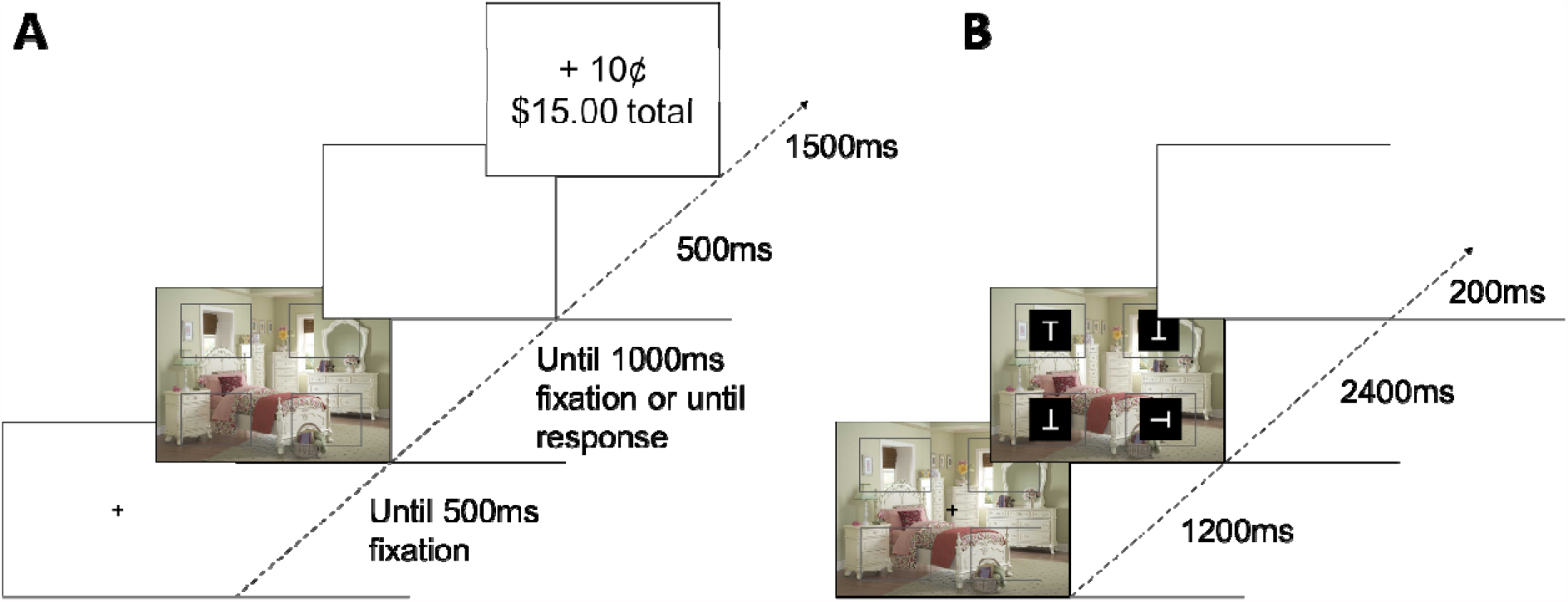
Time course of trial events during the training and test phases of the experiment. During the training phase (A) participants were presented with scenes with an empty box in each quadrant and instructed to pick a box by looking directly at it Depending on their choice, participants earn either 10c or 2c on every trial. During the test phase (B) participants were tasked with searching for a side-ways “T” among upright and upside down “T” distractors. Scenes previously experienced during the training phase were used as the background and were irrelevant to the task. Note that the stimuli are not drawn to scale in the figure, and the background has been changed from black to white for display purposes.

Participants were instructed to fixate (“look directly at”) the cross to begin each trial, then to “pick a box and look directly at it”. Participants were also informed that they would earn money on each trial, and the amount earned would depend on which box they looked in. Participants were encouraged to maximize their earnings by picking good boxes but were otherwise not provided any explicit information about which boxes were good. There were four 80-trial runs of the training phase during the in-lab visit, and two runs of abridged training phases that only had 40 trials while in the scanner. There were eight practice trials before the in-lab training phases where participants earned 5¢ on each trial but were informed that money earned was for demonstration purposes only. Eight different scenes were used in the experiment, totaling 50 presentations of each scene over 400 trials. The scenes were taken from the CB Database (Sareen et al., 2016) and were used in previous studies of value-driven spatial attention (Anderson & Kim, 2018a, 2018b). For each scene, one quadrant (and the box it contained) was designated the high-value quadrant and yielded a 10¢ reward while picking any other boxes yielded a 2¢ reward.

Participants were assigned to one of four training conditions in alternating fashion, with each quadrant of each scene serving as the high-value quadrant in one of the four conditions. The order in which the scenes were presented to each participant was randomized. If eye-tracking was unable to be conducted in the scanner, participants instead used two two-button response pads to indicate their selection on each trial (one button per quadrant) and received some initial practice trials to learn the button mapping. To be eligible for the scanning session, participants needed to earn at least $10.00 during the training phase runs conducted in the lab, which was taken to indicate sufficiently robust learning of the scene-reward contingencies.

### Test Phase

Each trial began with the presentation of one of the scenes from training along with the 4 rectangular boxes for 1200ms, followed by the presentation of a 1.1º “T” stimulus in white against a black background centered within each of the boxes. One “T” was tilted either 90º to the left or right and served as the target, while the other three “T”s were either upright or upside down (randomly determined with the constraint that all three non-target “T”s could not be oriented in the same direction). The display remained on screen for 2400ms during which participants could enter their response. For a subset of participants, eye-tracking data was also collected during this time and eye-positions within a region extending 4.6 º x 3.4 º beyond the boundary of the rectangle for a continuous period of 100ms were counted as fixations. Each trial ended with a blank inter-trial-interval (ITI) which lasted 1200, 1800, 2400, 3000, or 3600ms (equally-often). The fixation cross reappeared 200ms for the last 200ms of the ITI to indicate to the participant that the next trial was about to begin. The test phase consisted of six runs of 80 trials each for a total of 480 trials, with each scene being presented a total of 60 times. During each run there were 16 trials where the “T” displays never appeared and the scene continued to stay on the screen for 2400ms (non-search trials). During the 64 search trials (containing “T” display) in each run, the target appeared in each box/quadrant of each scene equally-often (and thus target position was unbiased with respect to which quadrant previously served as the high-value quadrant). The target was titled 90º to the left and right equally-often for each scene. Trials were presented in a random order. At the end of each run, the accuracy on the 64 target present trials was displayed for six seconds to provide performance feedback.

During the in-lab visit, participants were instructed to press the “m” key with their right-hand index finger if the vertical line of the sideways “T” was on the left, denoting an arrow pointing to the right. If the vertical line of the sideways “T” was on the right, participants were instructed to press the “z” key with their left-hand index finger. To become familiar with the mapping, participants had 8 practice trials that included feedback displays that said “Correct!” or “Incorrect!” depending on their response, after which participants had four runs of 80 trials to reach 85% accuracy and be eligible for scanning. If participants reached 85% accuracy in one of these runs, they became eligible and moved on to the next task. During the scan-center visit, participants were instructed to indicate the orientation of the target with their right-hand index and middle finger on the button response pad.

### Procedure

The experiment consisted of a lab visit for 1hr followed by a scan-center visit on the following day. During the initial appointment, participants provided their consent, completed the MRI safety screening, and were screened for adequate performance on the behavioral task. The majority of scene-reward training took place during the lab visit. Each eligible participant underwent fMRI in a single 1.25hrs session that took place the following day. Participants completed one run of the training phase, three runs of the test phase, an anatomical scan, another run of the training phase, and lastly completed three more runs of the test phase. The abridged training phases were completed to re-instantiate the space-outcome associations to protect against possible extinction (e.g., Lee & Shomstein, 2014).

### Measurement of eye position

During the lab visit, head position was maintained using an adjustable chin rest including a bar upon which to rest the forehead (SR Research). Participants were given a short break between different runs of the task, during which they were allowed to reposition their head to maintain comfort. During the fMRI scan, head position was restricted using foam padding within the head coil, and eye-tracking was conducted using the reflection of the participant’s face on the mirror attached to the head coil. Participants were given short breaks in between runs where they were allowed to close their eyes but otherwise were encouraged to remain still. Eye position was calibrated prior to each run of trials using a 9-point calibration (Liao & Anderson, 2020a, 2020b; Liao et al., 2020) and was manually drift-corrected by the experimenter during the initial fixation display as necessary. Due to the difficulty of measuring eye position in the scanner environment, eye data could only be acquired for a subset of participants (*n* = 19) during the scan session.

### MRI data acquisition

MRI data was acquired with a Siemens 3-Tesla MAGNETOM Verio scanner and a 32-channel head coil at the Texas A&M Translational Imaging Center (TIC), College Station, TX. An anatomical scan was acquired using a T1-weighted magnetization prepared rapid gradient echo (MPRAGE) sequence (150 coronal slices, voxel size = 1mm isotropic, repetition time (TR) = 7.9ms, echo time (TE) = 3.65ms, flip angle = 8º). Whole-brain functional images were acquired using a T2*-weighted echo planar imaging (EPI) sequence (56 axial slices, TR = 600ms, TE = 29ms, flip angle = 52º, image matrix = 96 × 96, field of view = 240mm, slice thickness = 2.5mm with no gap), using the same parameters as Kim and Anderson (2019c,2020a, 2020b). Each EPI pulse sequence began with dummy pulses to allow the MR signal to reach steady state and concluded with an additional 6 sec blank epoch. Each of the 6 runs of the test phase lasted 8.1 mins during which 810 volumes were acquired.

### Behavioral data analyses

In the training phase, performance was categorized in terms of how many times the high-value quadrant was chosen per run, averaged over the scenes. A one-way analysis of variance (ANOVA) was conducted on the proportion of high-value choice for each run, followed by pair-wise comparison between the first and last run of the training phase. Only training phase data collected in-lab were analyzed. In the test phase, RT was recorded from the onset of the four items comprising the search array, and RTs exceeding 2.5 SD of the mean of their respective condition or faster than 150ms were trimmed (2.78%). If eye-tracking data were available, the proportion of first saccade towards the high-value quadrant was compared to chance (25%); in addition, on no-target trials (which amounted to a free-viewing situation), total fixation duration was also computed for each quadrant and the mean for the high-value quadrant was compared to the mean of a given low-value quadrant (mean across low-value quadrants divided by three, paralleling Anderson & Kim, 2018a, 2018b). Only correct responses were analyzed. The effect sizes *d* were also computed, but the data were not otherwise transformed. Data were analyzed using SPSS and MATLAB, and figures were generated in Python.

### MRI data analyses

#### Preprocessing

All preprocessing was conducted using the AFNI software package (Cox, 1996). Each EPI run for each participant was motion corrected using the last image prior to the anatomical scan as a reference. EPI images were then coregistered to the corresponding anatomical image for each participant. The images were then non-linearly warped to the Talairach brain (Talairach & Tournoux, 1998) using 3dQwarp. Finally, the EPI images were converted to percent signal change normalized to the mean of each run, and then spatially smoothed to a resulting 5mm full-width half-maximum using 3dBlurToFWHM.

#### Statistical analyses

All statistical analyses were performed using the AFNI software package. A general linear model (GLM) was performed on the test phase data and included the following regressors of interest: (1) valid trial, reward/target quadrant on the left, (2) valid trial, reward/target location on the right, (3) invalid trial, both reward and target location on the left, (4) invalid trial, both reward and target location on the right, (5) invalid trial, target location on the left and reward location on the right, (6) invalid trial, target location on the right and reward location on the left, and no-target trials with (7) reward location on the left and (8) reward location on the right. Each regressor of interest was modeled using sixteen finite impulse response functions (e.g., Kim & Anderson, 2019c, 2020a, 2020b) beginning at the onset of stimulus presentation, and drift in the scanner signal was modeled using nuisance regressors.

To compare the peak of the haemodynamic response, the peak β value for each task-based regressor from 3-6s post search display onset was extracted (Kim & Anderson, 2020a, 2020b) and submitted to a priori paired samples *t*-tests. Two paired samples *t*-tests were conducted on the peak beta weight estimates. We compared trials where the target appeared in the previously high-value quadrant (valid) versus where the target appeared contralateral to the previously high-value quadrant (invalid), separately for each of the two hemifields (i.e., the peak of regressor 1 vs. 5 and 2 vs. 6) (as in Anderson et al., 2014; Anderson, 2017; Kim & Anderson, 2020a, 2020b). We focused on invalid trials in which the target was in the opposite hemifield as the previously reward-associated quadrant in order to isolate trials of maximal spatial competition between the target and reward history. A third paired samples *t*-test was conducted comparing no-target trials in which the previously reward-associated location was on the left and right (i.e.., the peak for regressor 7 vs. 8). The results were corrected for multiple comparisons using the AFNI program 3dClustSim, with the smoothness of the data estimated using the ACF method (clusterwise α < 0.05, voxelwise p < 0.005).

### Data availability statement

Deidentified raw behavioral and fMRI data for the experiment will be made available in a publicly accessible repository upon notification of acceptance of the manuscript, a link to which will be provided here.

## RESULTS

### Behavior

During the training phase, participants were able to learn the reward association. The proportion of trials on which the high-value quadrant was selected in the final run was high (0.962; see Figure 2A) and averaged across all runs (0.784) was well above chance, *t*(33) = 19.35, *p* < 0.001, *d* = 3.32. Pairwise comparisons show that participants on average made more high-value choices on the last run compared to the first, *t*(33) = 15.1, *p* < 0.001, *d* = 2.59. During the test phase, accuracy was high (97.5%) and participants were faster to respond to valid trials compared to invalid trials, *t*(33) = 7.99, *p* < 0.001, *d* = 1.37 (see Figure 2B). For the 19 participants we had eye tracking data for, initial fixations were significantly biased towards the high-value quadrant (33.6%), *t*(18) = 7.68, *p* < 0.001, *d* = 1.76. On trials where the target and distractors were not presented, total fixation duration was higher on high-value quadrants (2992 ms) compared to low-value quadrants (2012 ms), *t*(18) = 4.33, *p* < 0.001, *d* = 0.99 and initial fixations were significantly biased towards the high-value quadrant (38.4%), *t*(18) = 3.84, *p* = 0.001, *d* = 0.88. The behavioral data on both target-present and no-target trials fully replicate Anderson and Kim (2018a, 2018b).

**Figure 2.**
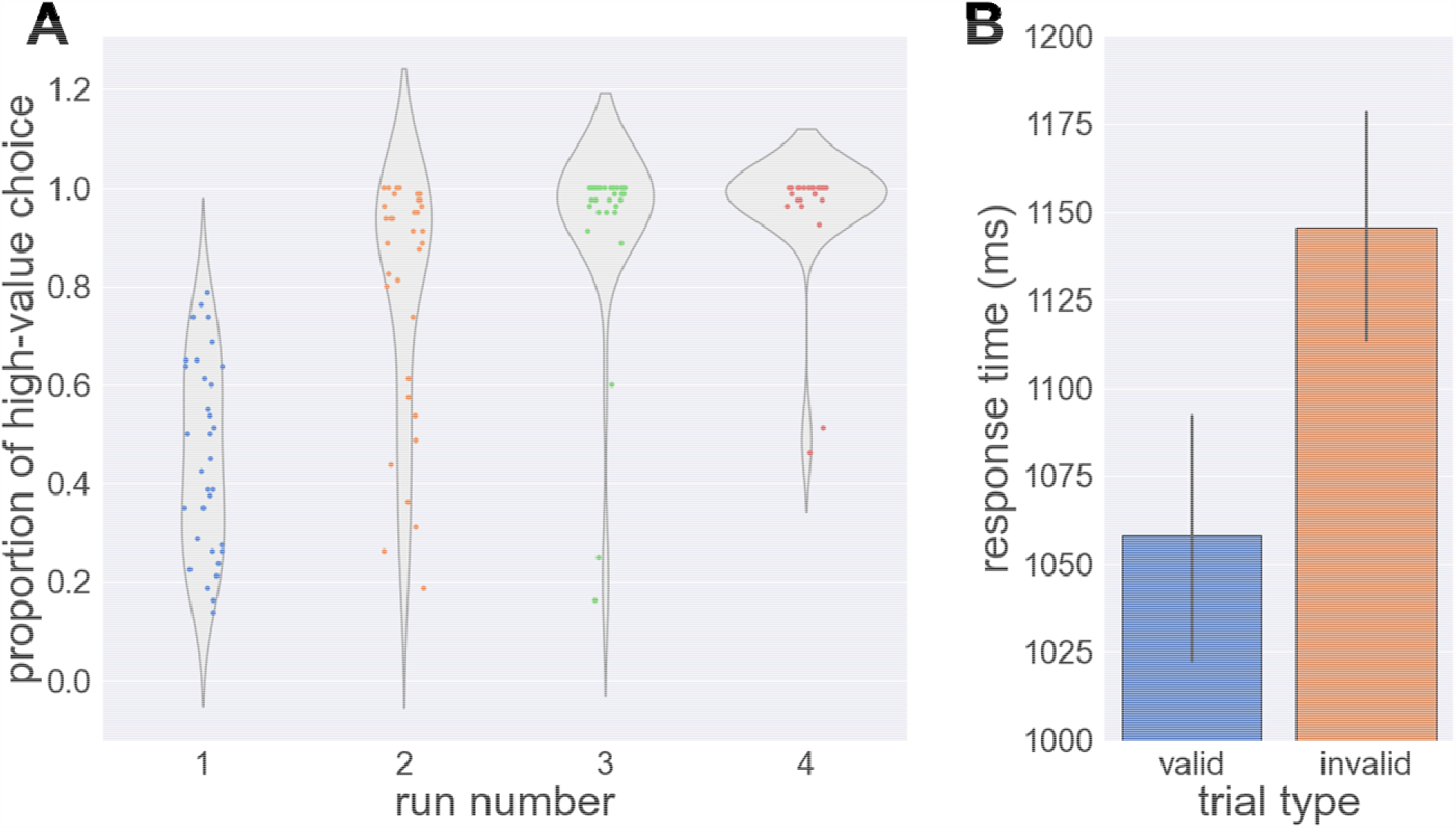
Behavioral results in the training and test phase. (A) Proportion of high-value choice by run during the training phase (B) Response time in the test phase by trial type. Error bars reflect standard error of the means.

### Neuroimaging

We compared valid trials on which the target appears in the same quadrant that was previously associated with reward against invalid trials when the target appears in a quadrant on the opposite hemifield (see Figure 3). Valid trials evoked elevated responses in oculomotor areas of the value-driven attention network (Anderson, 2017, 2019; Kim & Anderson, 2020a, 2020b) including the caudate tail, superior colliculus, and frontal eye field. We also observed increased activation on valid trials in the medial temporal lobes, particularly in regions associated with scene, space, and object processing like the hippocampus (O’keefe & Nadel, 1978), parahippocampal gyrus (Epstein & Kanwisher, 1998; Maguire et al., 1996), and the lateral occipital cortex (Grill-Spector et al., 2001). These are not regions typically associated with the value-driven attention network but may have been recruited to represent additional reward-related information in object-rich scenes. We also observed an increase in activity for the insula and anterior cingulate cortex (ACC), which have been previously implicated in reward-modulated attentional control (Hickey et al., 2010; Wang et al., 2015).

**Figure 3.**
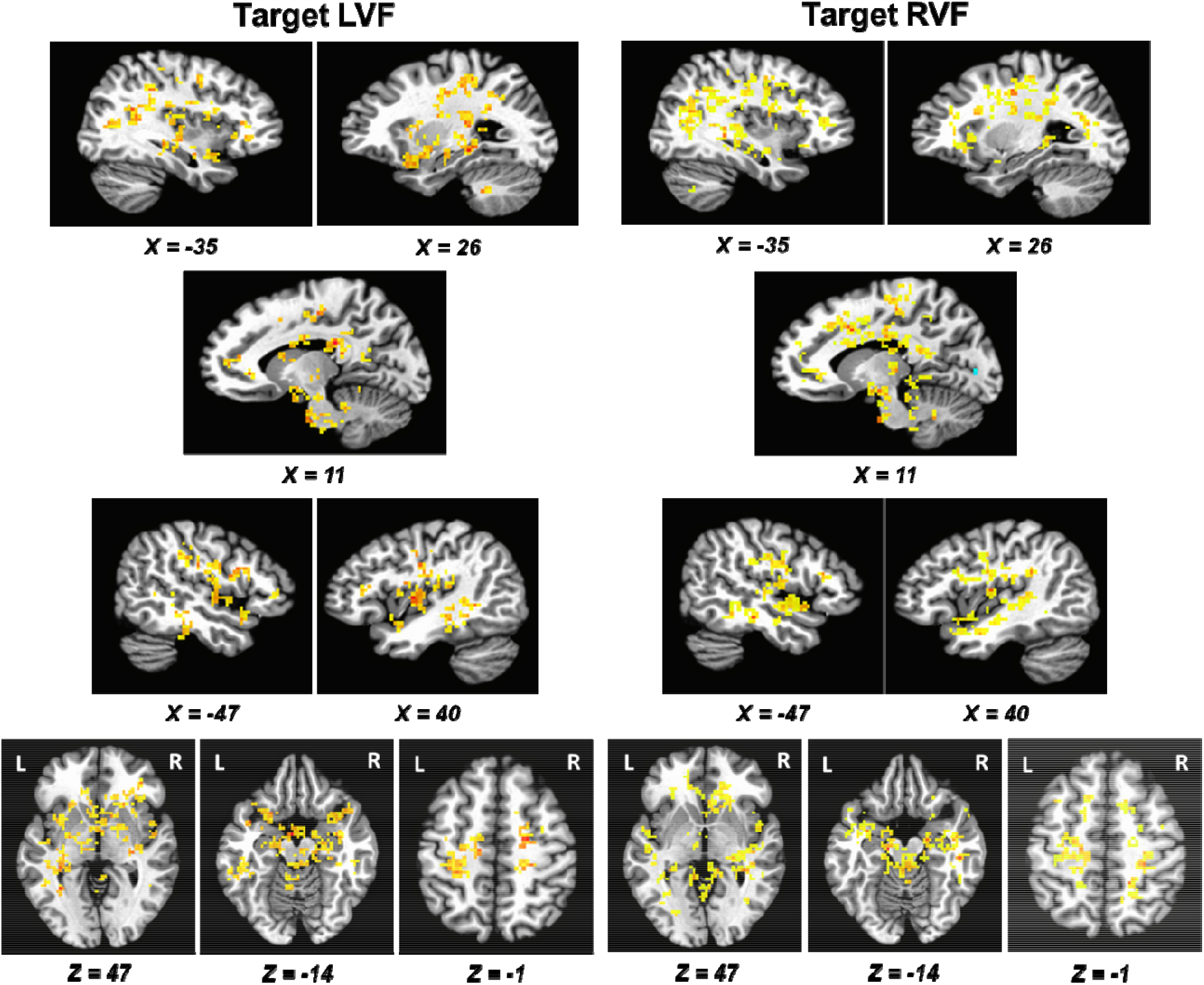
Regions that were significantly more active on valid compared to invalid (high-value quadrant in opposite hemifield) trials during the test phase for targets appearing in the A) right visual field (RVF) and B) left visual field (LVF). Activations are overlaid on an image of the Talairach brain. A complete list of all regions showing significant activation is provided in Supplemental Table 1.

We then probed no-target trials where only the scenes were presented, comparing trials where the previously reward-associated quadrant was on the left versus the right (see Figure 4). We observed increased activity in the right extrastriate visual cortex (see Figure 4) ipsilateral to the high-value quadrant. A complete list of all regions activated across all contrasts is provided in Supplemental Table 1.

**Figure 4.**
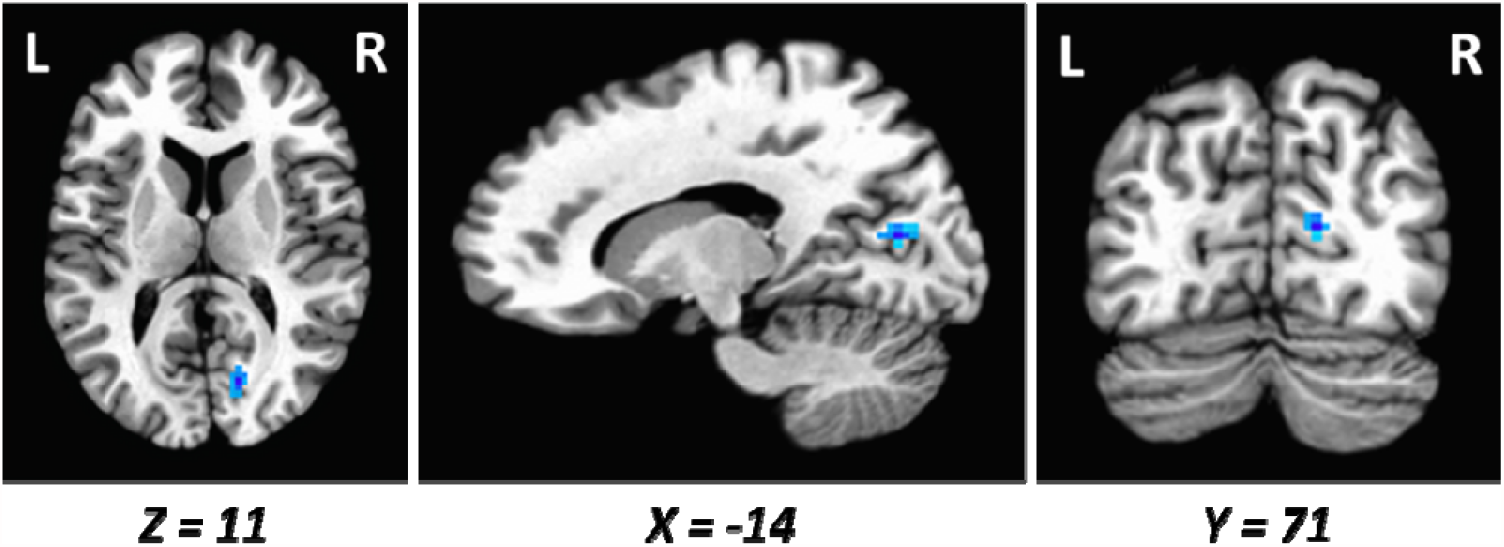
Regions that were significantly more active in response to the presentation of scenes without any targets or distractors during the test phase. The contrast depicted is the difference between previously high-value quadrant on the left and right (left – right) such that cooler colors correspond to stronger activations for previously high-value quadrant on the right (ipsilateral).

## DISCUSSION

In the present study, we investigated the neural basis of attentional priority for the region of a scene previously associated with reward. As in Anderson and Kim (2018a), we observed persistent spatial biases specific to different scenes in the form of cueing effects on RT and oculomotor biases; all behavioral measures indicated a persisting spatial bias towards previously high-reward locations in scenes. Our neuroimaging data comparing valid and invalid trials reveal some of the same neural correlates associated with value-driven feature-based attention, including the superior colliculus, frontal eye field, and caudate tail (Anderson et al., 2014, 2016, 2017; Anderson, 2017; Kim & Anderson, 2019b; Kim & Anderson 2020a, 202b; Hickey & Peelen, 2015). That is, targets evoked stronger activation in these regions when appearing in a previously reward-associated quadrant of the scene compared to when the previously reward-associated quadrant was in the opposite visual hemifield as the target. We also observed such elevated stimulus-evoked activity in the insula and ACC, each of which has likewise been linked to value-driven feature-based orienting (Hickey et al., 2010; Wang et al., 2015).

In this experimental paradigm, we used object-rich scenes to provide contextual information about the location of high-value quadrants. Accordingly, we observed increased activation on valid trials―in which the task required the participant to orient to the previously high-value quadrant―in regions of the brain known to play an important role in representing spatial layout, including the hippocampus, parahippocampal cortex, and occipito-temporal cortex (Epstein & Kanwisher, 1998; Maguire et al., 1996; O’keefe & Nadel, 1978). Such regions have not been previously implicated in value-driven attention and may be particular to reward’s modulatory influence on spatial orienting in scenes.

The caudate tail along with the superior colliculus and frontal eye field have been frequently linked to value-driven attentional capture by feature-defined stimuli (Anderson, 2016). Given its connections with the superior colliculus (Yamamoto et al., 2012) and its proximity to the medial temporal lobe (Segar, 2003), the caudate tail potentially serves a more general role in value-based attentional guidance by taking input from specific reward-associated features that are the targets of saccades as well as scene contexts associated with corresponding spatial priority. Such scene context representations may be mediated by the hippocampus, parahippocampal gyrus, and lateral occipital cortex (Epstein & Kanwisher, 1998; Maguire et al., 1996; O’keefe & Nadel, 1978). One of the most prominent spatial priority maps that guide attention is held in the posterior parietal cortex (Serences & Yantis, 2007; Sprague & Serences, 2013) and was not reliably activated in our task, in contrast to prior studies of feature-based value-driven attention (e.g., Anderson, 2017, 2019; Anderson et al., 2014; Kim & Anderson, 2020a, 2020b). The lack of parietal cortex activity may reflect a distinction between the representation of feature-based and scene-based spatial attention; the parietal cortex is situated closer to the occipital lobe and may be more suited for representing and prioritizing low-level features (Anderson, 2019; Serences & Yantis, 2007; Sprague & Serences, 2013) while information based on spatial context and layout is represented in the medial temporal lobe closer to the caudate tail (Epstein & Kanwisher, 1998; Maguire et al., 1996; O’keefe & Nadel, 1978). Our findings suggest an integrated neural system for value-based guidance with the caudate tail receiving priority computed from different regions in response to different objects or scene information and potentiating biased oculomotor behavior through the superior colliculus and frontal eye field.

On free-viewing trials where only the scene is presented, there were longer dwell times on previously reward-associated quadrants and these quadrants were more likely to be fixated first, replicating previous findings (Anderson & Kim, 2018a, 2018b). Such a bias in oculomotor behavior was associated with increased neural activity within the extrastriate visual cortex ipsilateral to the previously reward-associated quadrant. One interpretation of this finding is that by preferentially fixating on the previously high-value quadrant, there ends up being more visual information in, and thus stronger neural activity generated by, the hemifield opposite this quadrant once gaze has shifted. Therefore, this observed difference in neural activation on no-target trials may simply be a reflection of the oculomotor biases observed in behavior.

Some studies using a single high-value location against a blank background have shown that learned spatial biases do not persist into extinction (Jiang et al., 2015; Won & Leber 2016). Using scenes that comprised of objectless textures, Anderson and Kim (2018b) found a reliable spatial bias evident during free viewing but not during performance of a visual search task as in the present study (Anderson & Kim, 2018a). It is likely that the value-driven spatial biases observed in the present study require a distinct arrangement of objects within a scene, the spatial relationship among which serves as a contextual cue (Brockmole & Henderson, 2006a, 2006b). Accordingly, our observed neural correlates include visual regions traditionally associated with object processing like the lateral occipital cortex (Grill-Spector et al., 2001) and scene memory such as temporo-occipital and parahippocampal cortex (Epstein & Kanwisher, 1998; Maguire et al., 1996).

The ACC is often associated with control processes like filtering, resolving of conflict or gating of inputs (Mansouri et al., 2009), and is also thought to play a role in valuation signals that promote the repetition of a rewarded orienting response to a particular stimulus feature (reward-modulated priming; Hickey et al., 2010). In the present study, valid trials, which preferentially activated the ACC, also involve the repetition of a previously rewarded orienting response, in this case with respect to a spatial context. In this respect, our findings are further consistent with a parallel influence of reward on feature-based and spatial attention, with the recruitment of similar brain structures in spite of substantial differences between tasks.

In summary, our results suggest distinct neural correlates of value-driven spatial attention in the hippocampus, parahippocampal gyrus, and surrounding cortices, as well as core regions of a value-driven attention network that are recruited in support of both feature-based and spatial priority including the caudate tail, superior colliculus, and frontal eye field. Given its role in the control of eye movements (Yamamoto et al., 2012, 2013) and proximity to both the hippocampus and parahippocampal gyrus on the one hand and its connections with the ventral visual stream on the other hand (Seger, 2013), the caudate tail may be particularly suited to serve as a hub region playing a more central role in integrating value-based attentional priority across features and space, consistent with its central role in models of value-based attention (Anderson, 2019). We did not observe biased representation in the posterior parietal cortex, consistent with the dual mechanism of value-driven attention hypothesis (Anderson, 2019) and a distinctly feature-based mode of priority in this case. Our task incorporated object-rich scenes into the signaling of value, with scene-space reward contingencies, which may have resulted in scene and object-specific regions to be recruited into the value-driven attention network, highlighting a greater flexibility in the neural computation of value-based attention priority than previously assumed (Anderson, 2019). The novel correlates of value-driven attention observed here provide an impetus for future research investigating the role of the medial temporal lobe in creating context-specific value-driven attentional biases (see, e.g., Anderson, 2015).

## Acknowledgements

We thank Mark K Britton for assistance with data collection.

## Open Practices Statement

No experiment was preregistered. All anonymized study data, including the raw MRI data, and custom code will be available upon request. Data sharing for this article complies with the requirements of the funding agencies and the stipulations of the university IRB approvals.

