## Supplemental Table 1 for "Neural Correlates of Value-Driven Spatial Attention"

| A. **Validity Effect in LVF** | | |
| --- | --- | --- |
| **Region** | **x, y, z** | **Volume (ml)** |
| Left insula | 40, 6, 6 | 23.953 |
| Left putamen | 20, -11, 11 | 23.953 |
| Left parahippocampal gyrus | 16, 17, -16 | 23.953 |
| Left bed nucleus of stria terminalis | 23, -4, -11 | 23.953 |
| Left superior temporal gyrus | 38, -4, -13 | 23.953 |
| Right cingulate gyrus | -16, 27, 43 | 17.531 |
| Right frontal eye field | -20, 12, 47 | 17.531 |
| Right precuneus | -22, 46, 35 | 17.531 |
| Right middle occipital gyrus | -32, 66, 8 | 17.531 |
| Left hippocampus | 26, 31, -7 | 5.156 |
| Left caudate tail | 32, 33, -2 | 5.156 |
| Left lateral occipital complex | 49, 38, -9 | 5.156 |
| Left brainstem | 5, 26, -36 | 4.328 |
| Right postcentral gryus | -53, 12, 18 | 4.156 |
| Right inferior parietal lobule | -44, 29, 28 | 4.156 |
| Right caudate tail | -28, 21, -3 | 3.891 |
| Right hippocampus | -35, 29, -7 | 3.891 |
| Left frontal eye field | 21, 18, 44 | 2.703 |
| Right superior temporal gyrus | -40, -11, -21 | 1.984 |
| Right bed nucleus of stria terminalis | -44, -11, -11 | 1.984 |
| Left posterior cingulate gyrus | 11, 36, 21 | 1.859 |
| Left cerebellum | 6, 48, -16 | 1.797 |
| Left superior colliculus | 5, 31, -17 | 1.797 |
| Right superior colliculus | -2, 30, -12 | 1.797 |
| Left precentral gyrus | 34, 11, 41 | 1.172 |
| Left cerebellum | 14, 16, -31 | 1.156 |
| Left precuneus | 21, 51, 34 | 1.047 |
| Left temporoparietal junction | 51, 39, 15 | 0.984 |
| Right superior temporal gyrus (posterior) | -39, 24, 9 | 0.891 |
| Right lateral occipital complex | -48, 31, -12 | 0.828 |
| Right inferior frontal gyrus | -31, -24, -4 | 0.656 |
| Right precentral gyrus | -31, 1, 34 | 0.625 |
| Left cerebellum | -4, 54, -4 | 0.563 |
| Left anterior cingulate gyrus | 11, -31, 1 | 0.547 |
| Left posterior cingulate gyrus | 19, 54, 11 | 0.531 |
| Left claustrum | 21, -19, 4 | 0.516 |
| Left anterior cingulate gyrus | 11, -39, 14 | 0.422 |
| Left inferior frontal gyrus | 41, -29, 11 | 0.422 |
| Right medial frontal gyrus | -20, -41, 14 | 0.422 |
| Left cingulate gyrus | 14, -9, 41 | 0.422 |
| Left thalamus | 14, 21, 1 | 0.406 |
| Right thalamus | -1, 11, 4 | 0.375 |
| Left postcentral gyrus | 54, 16, 21 | 0.359 |
| Left postcentral gyrus | 54, 21, 34 | 0.359 |
| Left insula | 39, 31, 21 | 0.344 |
| Left postcentral gyrus | 61, 9, 14 | 0.328 |
| Right substantia nigra | -9, 20, -11 | 0.313 |
| Left parahippocampal gyrus | 31, 51, -1 | 0.313 |
| Right medial frontal gyrus | -11, -6, 49 | 0.313 |
| Left cerebellum | 26, 41, -36 | 0.297 |
| Left middle temporal gyrus | 41, 49, 1 | 0.297 |
| Left inferior parietal lobule | 39, 41, 29 | 0.297 |
| Right medial frontal gyrus | -11, -36, 36 | 0.281 |

| B. **Validity Effect in RVF** | | |
| --- | --- | --- |
| **Region** | **x, y, z** | **Volume (ml)** |
| Left parahippocampal gyrus | 13, 19, -14 | 52.563 |
| Right lateral occipital complex | -46, 26, -14 | 52.563 |
| Right parahippocampal gyrus | -20, 12, -14 | 52.563 |
| Right superior temporal gyrus | -44, 34, 8 | 52.563 |
| Right postcentral gyrus | -27, 33, 44 | 52.563 |
| Right frontal eye field | -22, 12, 44 | 52.563 |
| Right cingulate gyrus | -14, 31, 31 | 52.563 |
| Right posterior cingulate | -14, 41, 21 | 52.563 |
| Right caudate tail | -34, 37, -1 | 52.563 |
| Right medial temporal gyrus | -39, 43, 3 | 52.563 |
| Right superior colliculus | -4, 31, -5 | 52.563 |
| Left superior colliculus | 3, 34, -14 | 52.563 |
| Right hippocampus | -32, 17, -15 | 52.563 |
| Left substantia nigra | 6, 17, -12 | 52.563 |
| Right insula | -41, 18, 8 | 52.563 |
| Right middle occipital gyrus | -31, 61, 8 | 52.563 |
| Left frontal eye field | 21, 17, 47 | 22.953 |
| Left medial frontal gyrus | 11, 24, 57 | 22.953 |
| Left cingulate gyrus | 11, -11, 36 | 22.953 |
| Left cingulate gyrus | 11, 16, 28 | 22.953 |
| Left posterior cingulate | 11, 41, 18 | 22.953 |
| Left insula | 38, 21, 21 | 22.953 |
| Left insula | 31, 34, 21 | 22.953 |
| Left caudate tail | 21, 34, 11 | 22.953 |
| Left cuneus | 24, 75, 14 | 5.031 |
| Left posterior cingulate | 24, 62, 18 | 5.031 |
| Left superior temporal gyrus | 44, 32, 2 | 5.031 |
| Left caudate tail | 31, 41, 3 | 5.031 |
| Left hippocampus | 24, 37, 1 | 5.031 |
| Left middle occipital gyrus | 34, 59, 5 | 5.031 |
| Left parahippocampal gyrus | 30, 52, 8 | 5.031 |
| Right nucleus accumbens | -16, -14, -4 | 4.703 |
| Right insula (anterior) | -29, -19, 3 | 4.703 |
| Left anterior cingulate | 14, -40, 1 | 4.703 |
| Right claustrum | -26, -20, 14 | 1.844 |
| Left superior temporal gyrus | 46, 6, -4 | 1.203 |
| Left parahippocampal gyrus | 31, 4, -14 | 1.047 |
| Left fusiform gyrus | 41, 4, -19 | 1.047 |
| Right middle frontal gyrus | -39, -19, 19 | 0.984 |
| Left inferior frontal gyrus | 39, -11, -9 | 0.906 |
| Left lingual gyrus | 5, 76, 4 | 0.875 |
| Left paracentral lobule | -11, 29, 59 | 0.781 |
| Left medial frontal gyrus | 16, -41, 16 | 0.594 |
| Left inferior frontal gyrus | 26, -24, -6 | 0.547 |
| Left thalamus | 11, 24, 4 | 0.516 |
| Left anterior cingulate | 9, -16, 21 | 0.469 |
| Right precuneus | -24, 64, 24 | 0.469 |
| Right medial frontal gyrus | -16, -39, 21 | 0.406 |
| Right bed nucleus of stria terminalis | -16, -4, -11 | 0.375 |
| Left paracentral lobule | 9, 34, 66 | 0.359 |
| Left claustrum | 29, -1, 16 | 0.344 |
| Left supramarginal gyrus | 34, 49, 26 | 0.344 |
| Left superior temporal gyrus | 31, -11, -21 | 0.328 |
| Left thalamus | 1, 6, 1 | 0.328 |
| Right middle frontal gyrus | -29, -16, 31 | 0.328 |
| Left insula | 41, 11, 11 | 0.281 |

| C. **Free-Viewing Contrast LVF-RVF** | | |
| --- | --- | --- |
| **Region** | **x, y, z** | **Volume (ml)** |
| Right cuneus | -14, 72, 11 | 0.328 |

*Table 1 – All regions of the brain demonstrating significantly greater activations across all our contrasts. 1A and 1B contain the list of regions that had greater activation in response to valid compared to invalid trials. Valid trials were those where the target appeared in the previously high-value quadrant for that specific scene, while invalid trials were those where the target appeared in the opposite hemifield to the previously high-value quadrant for that specific scene. 1C contains the region that was more activated on free viewing trials for previously high-value quadrant on the RVF compared to the LVF. RVF = visual field; LVF = left visual field. x, y, z refer to the Talairach coordinates of the peak voxel of the cluster. Regions with the same volume of activation formed one contiguous cluster.*
